# Improving Bacterial Ribosome Profiling Data Quality

**DOI:** 10.1101/863266

**Authors:** Alina Glaub, Christopher Huptas, Klaus Neuhaus, Zachary Ardern

**Affiliations:** Chair for Microbial Ecology, Technical University of Munich, Freising, Germany; Core Facility Microbiome/NGS, Institute for Food & Health, Technical University of Munich, Freising, Germany

## Abstract

Ribosome profiling (RIBO-seq) in prokaryotes has the potential to facilitate accurate detection of translation initiation sites, to increase understanding of translational dynamics, and has already allowed detection of many unannotated genes. However, protocols for ribosome profiling and corresponding data analysis are not yet standardized. To better understand the influencing factors, we analysed 48 ribosome profiling samples from 9 studies on *E. coli* K12 grown in LB medium. We particularly investigated the size selection step in each experiment since the selection for ribosome-protected footprints (RPFs) has been performed at various read lengths. We suggest choosing a size range between 22-30 nucleotides in order to obtain protein-coding fragments. In order to use RIBO-seq data for improving gene annotation of weakly expressed genes, the total amount of reads mapping to protein-coding sequences and not rRNA or tRNA is important, but no consensus about the appropriate sequencing depth has been reached. Again, this causes significant variation between studies. Our analysis suggests that 20 million non rRNA/tRNA mapping reads are required for global detection of translated annotated genes. Further, we highlight the influence of drug induced ribosome stalling, causing bias at translation start sites. Drug induced stalling may be especially useful for detecting weakly expressed genes. These suggestions should improve both gene detection and the comparability of resulting ribosome profiling datasets.

## Introduction

Ribosome profiling is a recently established method of RNA sequencing and conceptually straightforward. Translating ribosomes of a bacterial culture are isolated and treated with RNases *in vitro*. The ribosomes protect mRNA fragments – the ‘footprints’, which are then isolated and sequenced in high-throughput. This allows taking a snapshot of the ongoing translation at the time point of cell harvest, hence the ‘translatome’ is observed in contrast to the transcriptome obtained in common RNA sequencing [1–3]. Important experimental steps include RNA digestion, monosome purification, RNA extraction and footprint size selection [1, 4] (Figure 1). However, each of these steps can be performed using a number of different variations, all of which influence the final data. In the following, each critical step and available methods are introduced in some detail.

**Figure 1:**
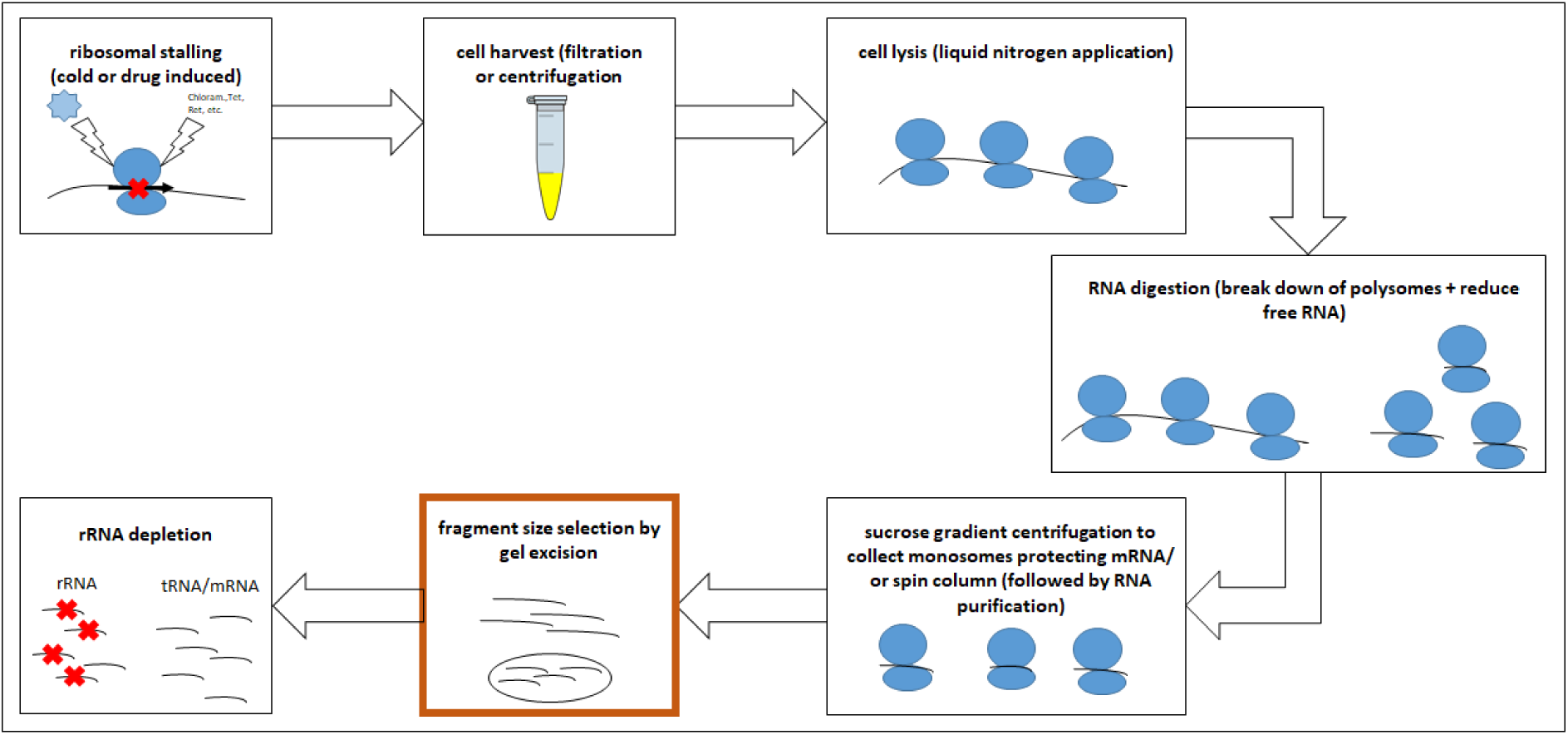
Overview of crucial experimental steps having an influence on the RNA fragments obtained for ribosome profiling. The step we primarily want to focus on is highlighted.

### Ribosome stalling in general and for start site detection

Since ribosomes are harvested from living cells, in order to observe all actively translated regions we want to obtain a single snapshot in time. By stalling the ribosomes, potential adaptation to environmental changes in the translatome due to subsequent processing steps is minimized [1, 5], and only mRNA actively translated at the time of harvest is obtained for sequencing. Ribosome stalling is induced either through application of drugs or rapid cooling [1, 4].

Chloramphenicol has been the most commonly used antibiotic for inducing stalling thus far; however, recent studies have shown that it does not fully inhibit prokaryotic translation. While chloramphenicol stops the elongation of most ribosomes, initiation still progresses, which leads to an accumulation of ribosomes at the start codon [6]. To avoid this bias, rapid filtration for harvest combined with quick liquid nitrogen freezing can be used instead [6, 7]. With this method no ribosomal accumulation at the translational start site is observed [6, 8]. Three studies included in our survey used filtration and flash freezing [9–11].

In contrast to the unbiased ribosomal stalling which is desirable in certain contexts, some studies intentionally use a drug-related bias for translation start sites precisely because of the improved detection of a gene’s start codon. For instance, in bacteria, tetracycline stops protein elongation by preventing tRNA binding to the ribosomal A-site due to its direct overlap of the anticodon [12]. Thus, tetracycline has been used for start site mapping. However, as this blocking is a reversible process, the trapping of the start site is only semi-specific [12, 13]. A study performed by Meydan et al. (2019) tested the antibiotic retapamulin (Ret) from the class of pleuromutilins as an elongation blocker [14]. Another drug applied for start-site mapping is the peptide Onc112, which was shown by Weaver et al. (2019) to be suitable in stalling ribosomes precisely at the initiation site by destabilizing the initiation complex such that subsequent elongation is prevented [15]. Although Ret and Onc112 show differences in the distance to an upstream start codon, both allow precise start site detection [16].

### Cell harvesting and lysis

Mainly two methods are used for harvesting, either rapid filtration or centrifugation. For rapid filtration, the culture is quickly transferred to a membrane and filtered by applying vacuum pressure [7]. Cells are scraped from the membrane and flash-frozen in liquid nitrogen. The alternative is centrifugation of cooled cell suspensions [8, 17]. For both, proceeding quickly is necessary to guarantee ribosomal stalling before any cold-shock response alters the translatome [17, 18]. Rapid filtration is the method of choice as it seems to result in less variation, possibly due to faster and hence better ribosomal stalling [5, 11, 17].

Before freezing the cells in liquid nitrogen, lysis buffer should be added [5, 17]. The buffers used vary in their composition but should guarantee the stabilization of the ribosome-bound mRNA complex [17]. Negative effects may occur if excess amounts of some salts such as magnesium are used [8, 17, 19, 20]. These either destabilize ribosomes thereby releasing the mRNA or conversely strengthen folding of unbound mRNAs, preventing digestion and leading to these fragments being mistaken as footprints [17]. The frozen cells are pulverized in a grinder mill or using a mortar and pestle to release the ribosomes [8, 17, 21].

### Ribosomal footprint generation

The prepared ribosomes are incubated with endo- and/or exonucleases to digest any unprotected mRNA, aiming at monosomal ribosomes with a footprint inside [4, 17]. Polysomes will dissolve to monosomes as unprotected mRNA is digested [4, 17]. The length of remaining protected fragments inside the ribosomes is given by the size of the ribosome. For eukaryotes, the ribosomal footprints are about ~ 28 - 30 nucleotides [2, 4, 5]. In contrast, the ribosome footprint length is still highly debated for prokaryotes. Some studies hypothesize a length of 23 or 24 nucleotides due to this being the most common fragment length [6, 22, 23]. In contrast, other studies including a few of those examined here use ‘ribosomal footprint’ in a much larger range of up to 42 nucleotides [9, 10, 24].

Now the question of the RNases useful for ribosome footprinting arises. In eukaryotes, the endonuclease RNase I is used in many studies. RNase I has no cleaving bias for specific nucleotides and the unprotected mRNA seems to be trimmed right to the edge of the ribosome [4, 5]. Interestingly, RNase I is claimed to not work in bacteria and especially *E. coli*, as it is bound by the 30S ribosome subunit and therefore inhibited [6, 25]. Nevertheless, ribosome profiling experiments have been conducted successfully using only RNase I [26, 27], or in combination with other enzymes [28] producing footprints of a size of ~23 nt. Due to the assumption of RNAse I unsuitability, the enzyme micrococcal nuclease (MNase) is normally used for prokaryotic experiments instead, despite this enzyme having some sequence specificity [29], resulting in greater variability in ribosome footprint lengths.

In eukaryotes, it is possible to detect reading frames for single proteins. The mRNA progresses stepwise through the ribosome, showing a periodicity of three, i.e. codon-wise. Thus, a single-codon resolution is achievable [2, 6]. In contrast for prokaryotes, where reading frames can be detected in a sum signal, the resolution of standard ribosome footprinting methods lack precision for reading frame determination in individual genes [30]. An approach for increased precision uses RelE in addition to MNase. RelE only cleaves the mRNA in the A-site of the ribosome if activated in a stress condition, and normally cleaves between the 2^nd^ and 3^rd^ base of a codon [31–33]. Therefore RelE is suitable for reading frame determination [31]. Since RelE cleaves the footprint within the ribosomes, in addition to MNase cleaving the unprotected mRNA outside the ribosomes [34, 35], the resulting fragments are even shorter, which can be harder to map accurately [31]. Consequently, careful size selection may be particularly important if using RelE.

### Performing size selection and rRNA depletion

To enrich for ribosomes from the crude cell lysate after RNase digestion, sucrose gradient centrifugation is performed [5]. There are, however, limitations in detecting and obtaining the layer containing the monosomes. Therefore, enrichment of ribosomes can also be performed using gel filtration [4, 36]. Nevertheless, gradient centrifugation is still the most common method [11, 31, 37, 38]. From the enriched ribosome fraction, the total RNA is isolated. Putative ribosomal footprints are separated from other RNAs (e.g. rRNA and tRNA) by performing a size selection using gel electrophoresis. Samples are loaded onto, e.g. a TBE-urea gel and certain fragment sizes are excised after comparison to a corresponding DNA or RNA ladder [5, 30]. Another possibility for fragment size selection is to use a size exclusion spin-column, but this has rarely been used [39]. Size selection is a crucial step since it seems to have a major impact on the sequencing results. However, up until now, there is no consensus about which read lengths to choose when conducting bacterial ribosome profiling. In the beginning of ribosome footprinting of bacteria several groups isolated reads ranging from 28 to 40 nt [11, 40], while others chose fragments of 23±3 nt [23], and some groups started with fragments as short as 15 nucleotides [41, 42]. In a recent study, Mohammad et al. (2019) claimed that the diversity of bacterial ribosomal footprints is dependent on the characteristics of prokaryotic ribosomes and therefore postulate that a broad range of read lengths (15 to 40 nucleotides) should be taken for analysis. They claim that sampling this full range of read lengths is the most useful approach as it yields the most informative output, with a peak at around 24 nucleotides [6, 22]. Several studies have already used this range [6, 41, 42]. Unfortunately, the RNases used also degrade the ribosomal rRNA to some extent. Therefore, rRNA depletion is conducted, helping to ensure that more mRNA-reads are sequenced. Ingolia provided a method for depletion in the first RIBO-seq protocol published [5], but since then different kits have become available for prokaryotic rRNA removal, such as RiboZero (Illumina), RiboMinus (ThermoFisher), MICROBExpress (ThermoFisher) and others. There are publications available comparing the efficacy of rRNA removal for the different kits, thus this is not discussed further [43, 44].

### Evaluation of RIBO-seq data

To discriminate between untranslated and translated RNA, i.e. between non-coding RNA and mRNA, a direct comparison between RIBO-seq and RNA-seq results of the same culture can be made. Evidence for translation of open reading frames (ORFs) is given by dividing the reads per kilobase per million (RPKM) values obtained in RIBO-seq by the values from RNA-seq experiments. This ratio, designated ribosomal coverage value (RCV), is a measure of the extent to which an mRNA is translated. It has been suggested that an ORF should be considered as expressing a protein if the RCV is at least 0.355 [27], although the appropriate threshold will vary somewhat between strains and samples.

The diversity of methods that have been employed and the increasing use of RIBO-seq prompted us to examine and compare available bacterial ribosome profiling data sets to assess the methods used and their influence on the output. As other studies have already investigated the influence of stalling and the RNases used for ribosome footprinting, we particularly focus on the size selection performed and the resulting read length distribution to shed light on its influence on the final outcome. Further, we would like to make recommendations for future prokaryotic experiments.

## Methods

### Sample selection

For our analysis, we compared 48 available RIBO-seq *E. coli* K12 samples from 9 different experiments (Table S1) [9–11, 24, 31, 37, 38, 45, 46], with all samples grown in lysogeny broth (LB) medium. Only substrains of K12, i.e. BW25113/BWDK, MG1655 and MC4100 were considered here because of their close phylogenetic relationship (23 samples aligned to MG1655, 15 samples aligned to BW25113, 10 samples aligned to MC4100) [47–50]. Besides this, more importantly there are differences in the experimental procedures for stalling and harvesting the cells, and the size selection performed, allowing observations about their influence on the outcome. Abbreviations of each dataset were created for further use (Figure 2). Original GEO numbers, study abbreviations and the experimental variations are listed in Table S1. Of particular interest to us is the effect of the size selection step on the read length distributions of trimmed and mapped reads. One important scientific question related to this concerns whether a particular length is most suitable for detecting protein-coding ORFs.

**Figure 2:**
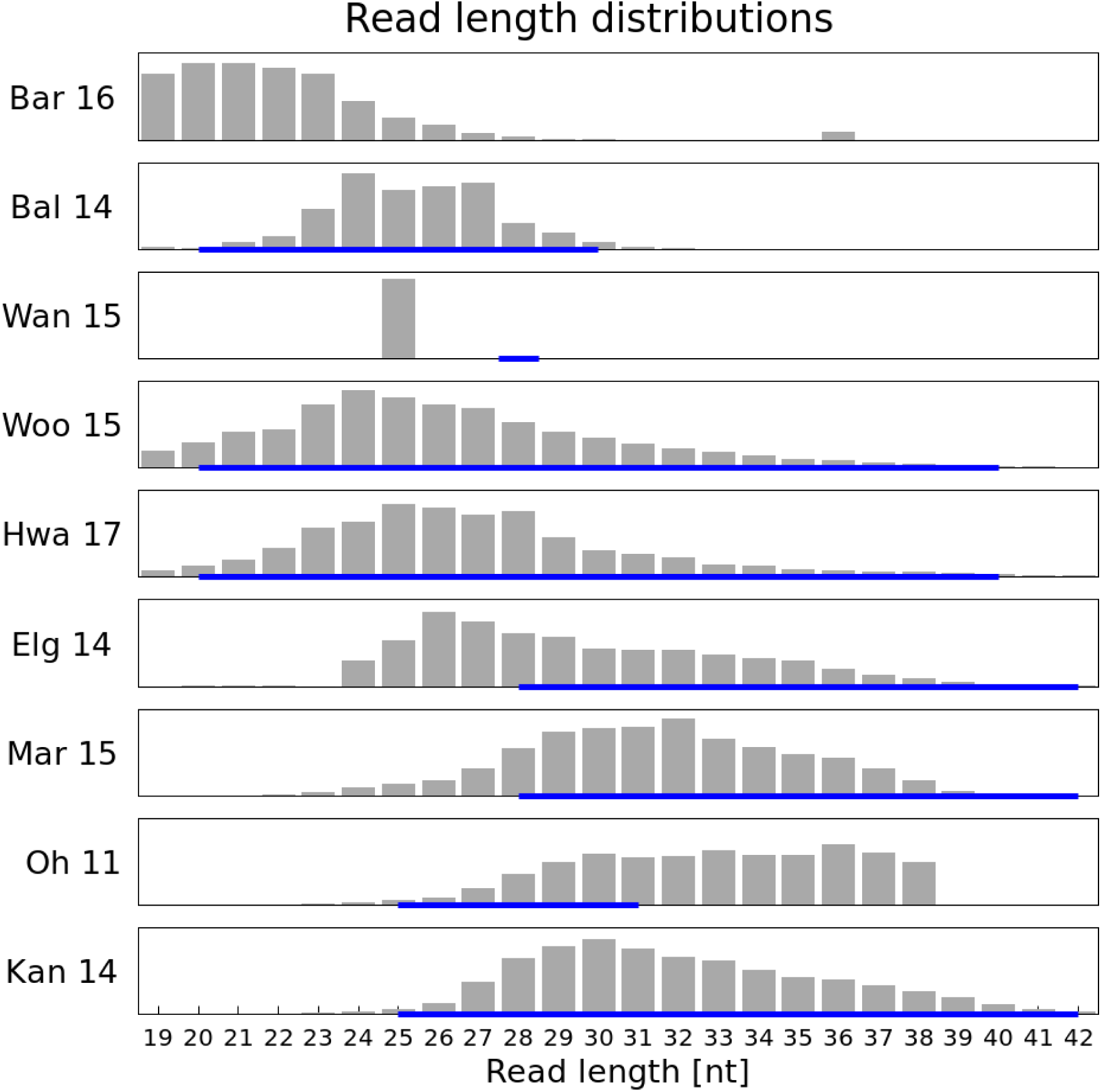
Read length distribution of different samples (one example per project) with reported size selection (blue line below) after trimming

### Bioinformatic data analysis

Our analysis pipeline started with inspection of raw fastq files using FastQC v0.11.4 (http://www.bioinformatics.babraham.ac.uk/projects/fastqc/). From this, the observed adapter contaminations and over-represented sequences can be inferred. Before performing the analysis, another frequently used adapter (5’-CTGTAGGCACCATCAAT-3’) [11, 38, 45] was added to the existing FastQC adapter list. Two experiments each used unique adapters, which were defined as input settings for the trimming [10, 46]. For the remaining four experiments the adapter sequences used were not stated [9, 24, 31, 37]. All samples were inspected using fastqc and adapters or other contaminants (overrepresented sequences) were identified. FastQC is only able to detect adapter sequences specified in the program. If available we used the published or detected adapter for trimming, otherwise we chose the most overrepresented sequences.

The software used during processing data are summarized in Table S2, with settings given if differing from default. Bash scripts used are available upon request. Trimming was performed using fastp version 0.14.2 [51]. Firstly, the identified adapter sequence was trimmed, and secondly, if present, overrepresented sequences above three percent were chosen for trimming as well. For some samples from the set Bal_14, for example, a second trimming step was performed, as a single overrepresented sequence constituted over 30% of the remaining reads. A BLAST analysis of this sequence identified it as rRNA fragment, which could also have been filtered out later in the pipeline. Automatic detection of adapters by fastp, followed by their trimming, was not successful even though an automatic trimming should be possible according to the program description. Therefore, we provided the sequences for trimming. Subsequent mapping of reads was performed using Bowtie2 version 2.2.6 with local alignment [52]. Reads mapping to rRNA or tRNA were moved into a new file using a custom bash script. Prediction of translated open reading frames was performed with REPARATION, a ribosomal profiling assisted reannotation tool [53]. The REPARATION workflow was adjusted by replacing UBLAST with DIAMOND for choosing the training set, after ORF prediction with prodigal. Predictions matching annotated genes were then used to analyse the total amount of mappable reads in a sample (after removal of rRNA/tRNA) necessary for detection of the genes. For a second verification of the translation status of these genes, the ribosomal coverage value (RCV) was calculated. Besides ribosome profiling, RNA sequencing data is also necessary for this evaluation, and was available for 22 of our 48 chosen samples.

Trimmed reads mapping to annotated protein-coding mRNA were analysed regarding their length distribution. The resulting length distributions were compared to the published experimental size selection used. Reads mapping to either tRNA or rRNA were also analysed regarding their length variation, testing whether different read lengths correlate with specific types of RNA. Read length distribution analyses were performed with custom bash scripts. We also investigated potential differences in read lengths mapping upstream of or directly within the start region of genes. The length distribution patterns in either the start region (from the start codon of an annotated gene to 25 nucleotides downstream of the start) or in the 5’ untranslated region (5’-UTR; in our case 25 nucleotides upstream of the start codon of each annotated gene) for all annotated genes were calculated. Additionally, the stop region read length distribution (25 nucleotides upstream of a stop codon) and the distribution for the remaining gene (between start+25 and stop-25) were calculated. Similarly, the median amount of reads, based on the read distribution in the region of interest, was calculated for all annotated genes per sample.

Claims that chloramphenicol induces ribosomal stalling and causes a read accumulation at the start region were also investigated. All samples from set Oh_11 were used here, as four were treated with chloramphenicol whereas the remaining four were not. However, harvesting methods also differed between the two sample subsets; chloramphenicol samples were centrifuged whereas untreated samples were harvested by rapid filtration. Nevertheless, in this set of eight samples, annotated genes were analysed regarding their ribosome footprint RPKM values and categorized for further analyses of high, medium, and weakly expressed genes (Table S3). For each category, ten genes were chosen which showed approximately the same expression status throughout all eight samples. P-site locations were inferred as 15 nucleotides upstream of the 3’ end of each read. By counting this specific position, each read will only be considered once for further analysis. This way the p-site locations of each read in the region starting from the translation start site and continuing for an additional 50 nucleotides were counted in order to investigate potential accumulation of reads at the start position. For each category, mean values at each unique position were calculated and compared between treated and untreated samples, both normalized. In addition, a similar comparison based on the median read amount at each unique position per subset was performed to verify the obtained results (data not shown).

## Results

### Size selection

For eight experiments, we were able to find information about the gel size selection performed (Table S1). We were interested in reads between 19 to 42 nucleotides, which should represent footprints. However, for several samples (from sets Oh_11, Woo_15, Mar_16, Kan_14, Hwa_17, Bal_14) reads longer than 42 (not shown) were detected, possibly due to insufficient trimming. These were not taken into consideration for the read length distribution analysis, as their range is located outside our range of interest, i.e. they might not represent ribosomal footprints. In general, analysing the trimmed and aligned reads mapping to mRNA and omitting reads mapping to either rRNA or tRNA, we expect to find a footprint distribution pattern matching the reported size selection thresholds in each publication. In reality, we find that for some samples the distribution patterns differ significantly from the reported size selection. Figure 2 shows the actual distributions for one representative sample per experiment.

For set Bar_16, we were not able to find any information about their performed size selection during gel excision. However, the two samples belonging to this experiment showed a peak value at 20 to 21 nucleotides, i.e. had the shortest fragments present in the samples (Figure 2 first panel). The third panel shows the discrepancy for experiment Wan_15 between a performed size selection of around 28 nucleotides and the actual read length peaking at a length of 25 nucleotides. For these samples, adapter trimming had already been performed for the data released by the group. In set Elg_14 (Figure 2 sixth panel), size selection was performed between 28 to 42 nucleotides. The actual distribution starts at 24 nucleotides and has a peak value at 26 nucleotides. Interestingly, this peak should have been excluded by their size selection.

For the sample Oh_11 (Figure 2 eight panel), a size selection from 25 to 31 nucleotides was performed. The distribution starts at around 27 nucleotides up to a length of 38, covering most of but also exceeding the intended range. In set Kan_14 (ninth panel), a size selection between 25 to 42 nucleotides was performed resulting in a peak at a length of 30 nucleotides. For the remaining two samples Woo_15 and Hwa_17 (Figure 2 fourth, fifth panel), the highest peak can be seen at either 24 or 25 nucleotides, matching the range of selection. With their performed selection, the resulting peak values are located in the lower half of the chosen range. For four experiments, namely Bal_14, Hwa_17, Kan_14, Woo_15, their actual read length distribution is situated within their performed size selection (Figure 2 second, fourth, fifth, eighth panel). The chosen ranges varied between 20 to 30 or 40 nucleotides with peak values between 20 and 30 nucleotides.

Results from this analysis imply that fragments of 24 to 27 nucleotides length are the most frequently protected. Therefore, they might be the most informative regarding gene positions. To test this hypothesis, we performed an analysis focusing on the length of reads aligned to annotated protein-coding genes compared to rRNA and tRNA genes.

### Specific read lengths correspond to different RNA types

To test whether different RNA types (mRNA, rRNA, tRNA) vary in their read length, we analysed the lengths between 20 to 40 nucleotides for each sample. The two samples from experiment Wan_15 were excluded from analysis, since the length distributions in these samples were narrow (Figure 2C). The percentage at the analysed read lengths for each RNA type was calculated, with subsequent calculation of medians (Figure 3). A peak at around 24-27 nucleotides was observed for mRNA, with tRNA and rRNA more likely to have longer reads, especially tRNA. Additional calculations were performed regarding the mean values for either the read length distributions within an RNA type or the mean value per type at a specific length and can be found in supporting information (Figure S1, S2).

**Figure 3:**
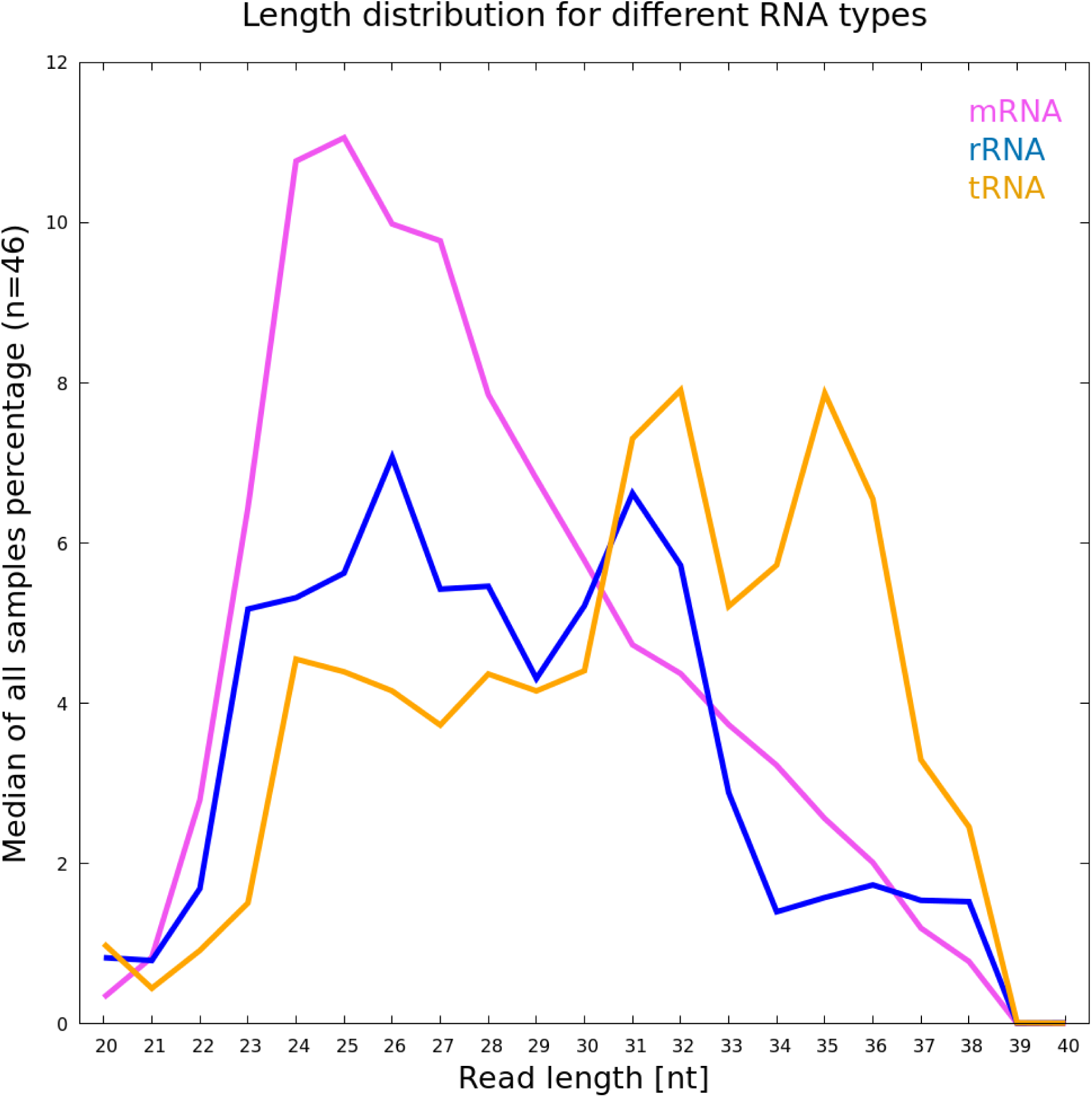
Read length analysis for three RNA categories depending on the median of percentage of RNA type calculated for each sample (n=46). Pink= mRNA, blue = rRNA, orange = tRNA

In our analysis we detect a peak at 24/25 nucleotides with a slightly lower amount of reads with a length of 26 to 27 nucleotides for mRNA (Figure 3, pink line). This supports the hypothesis that reads with these lengths might be the most informative ones regarding protein-coding genes. Thus, we suggest choosing a range between 22 to 30 nucleotides for size selection to obtain the most highly informative reads. For reads mapping to rRNA, a peak can be detected at 26 nucleotides of length (Figure 3, blue line). Located fully within our suggested range these reads cannot be excluded due to a narrower selection. Therefore, additional rRNA depletion during the experiment is advised to minimise the amount of reads that are wasted sequencing rRNA. The second peak of 31 nucleotides would be excluded with our suggested size selection. Reads mapping specifically to tRNA are predominantly longer, having two peak values at a length of 32 and 35 nucleotides (Figure 3, orange line). Again, with our suggested size selection these fragments could mostly be excluded.

### Longer reads in 5’-UTR region

A hypothesis that warrants further investigation claims that reads in the range of 28 to 40 nucleotides are associated with incorporated Shine-Dalgarno (SD) motifs [40, 41]. We thus speculate that reads mapping upstream of a start codon may also be longer due to SD sequences in this region. For this comparison, we chose to analyse the read range from 24 to 40 nucleotides. Thus, because of their narrow size selection ranging on average from 20 to 30, experiments Wan_15 and Bal_14 were excluded from this analysis. Further, Bar_16 was also excluded since no size selection was reported and the upper part of the examined range of reads was also missing. This left 30 samples for the analysis.

A clear trend for both of the analysed regions can be seen. Reads mapping directly in the start region tend to be shorter with 27 nucleotides being the most frequent length (Figure 4). However, reads mapping in the 5’-UTR region of genes are substantially longer, with 34 nucleotides as the most common length (Figure 4). The SD motif, oftentimes located in the upstream region [66], potentially results in longer reads due to the interaction between SD sequence and mRNA. Additionally, boxplots showing the amount (in percentage) of reads for each length over all samples were created (Figure S3A-D). Results for the analysis of the stop region and over the whole gene length are also included. The most prominent read length for these two regions again seems to be 27 nucleotides in length (Figure S3C, S3D). These results also support our hypothesis that this length is particularly associated with protein-coding characteristics.

**Figure 4:**
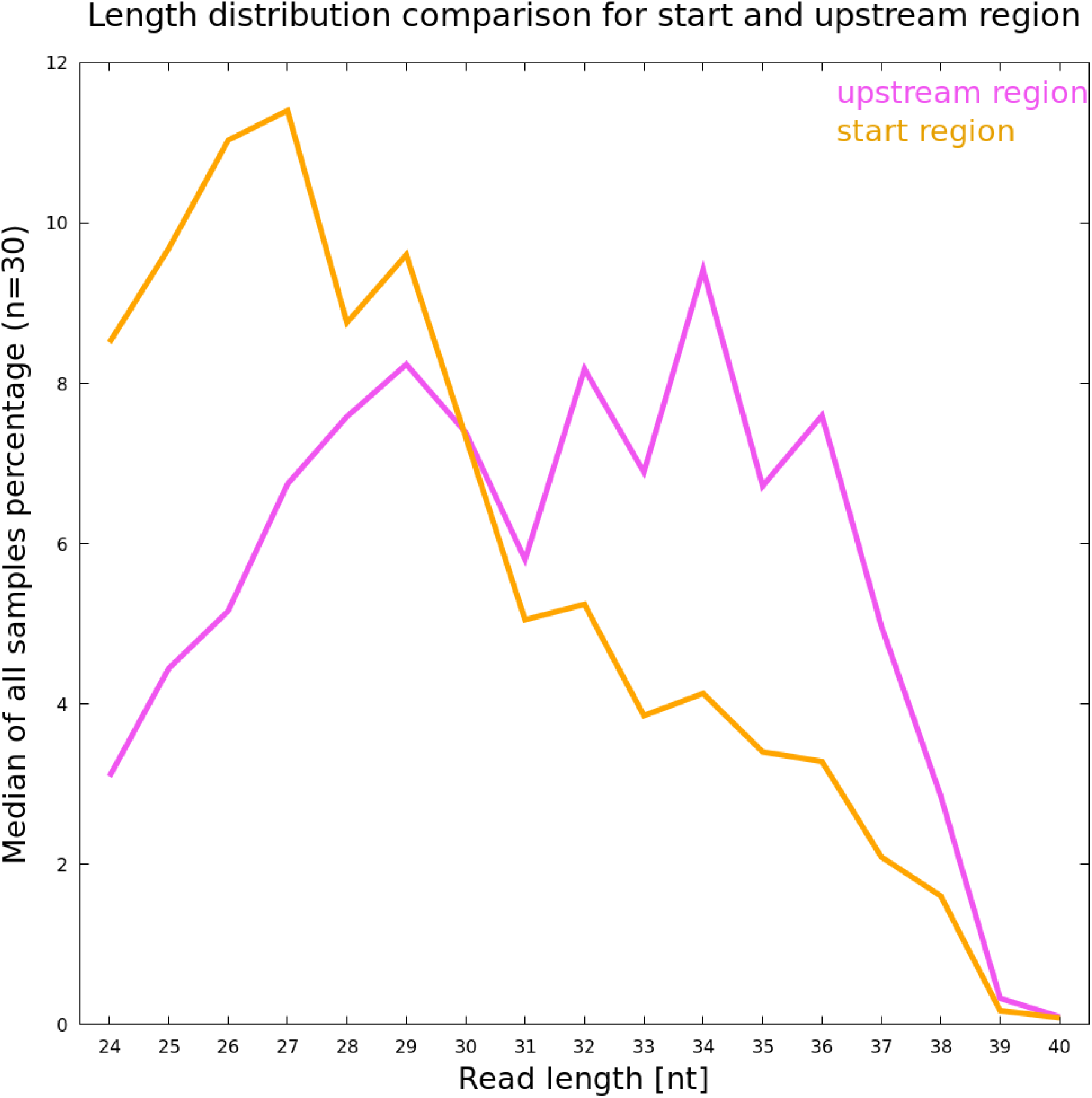
Prevalence of specific read lengths in the start and 5’-UTR region (potentially including Shine-Dalgarno sequence) based on their median

### Potential influence of chloramphenicol on read length

The hypothesis of drug induced stalling of ribosomes at the translation initiation site (TIS) was examined. We can confirm this phenomenon, i.e. ribosome footprints in the start region of genes are enriched in drug-treated samples. However, we see different results for genes expressed at different levels. Shown in Figure 5 is the average read accumulation in the TIS (translation start plus 50 nucleotides downstream). For highly expressed genes (Figure 5A), drug induced stalling results in an only slightly higher accumulation of reads at the TIS compared to untreated samples. Further translation results in several additional accumulations along the mRNA in treated samples. Drug application leads to a more clearly increased amount of reads located at the TIS for genes expressed at an intermediary level (Figure 5B). Interestingly, the largest effect of stalling is observed for weakly expressed genes (Figure 5C). This comparison suggests that the effect of chloramphenicol is dependent on the expression status of genes.

**Figure 5:**
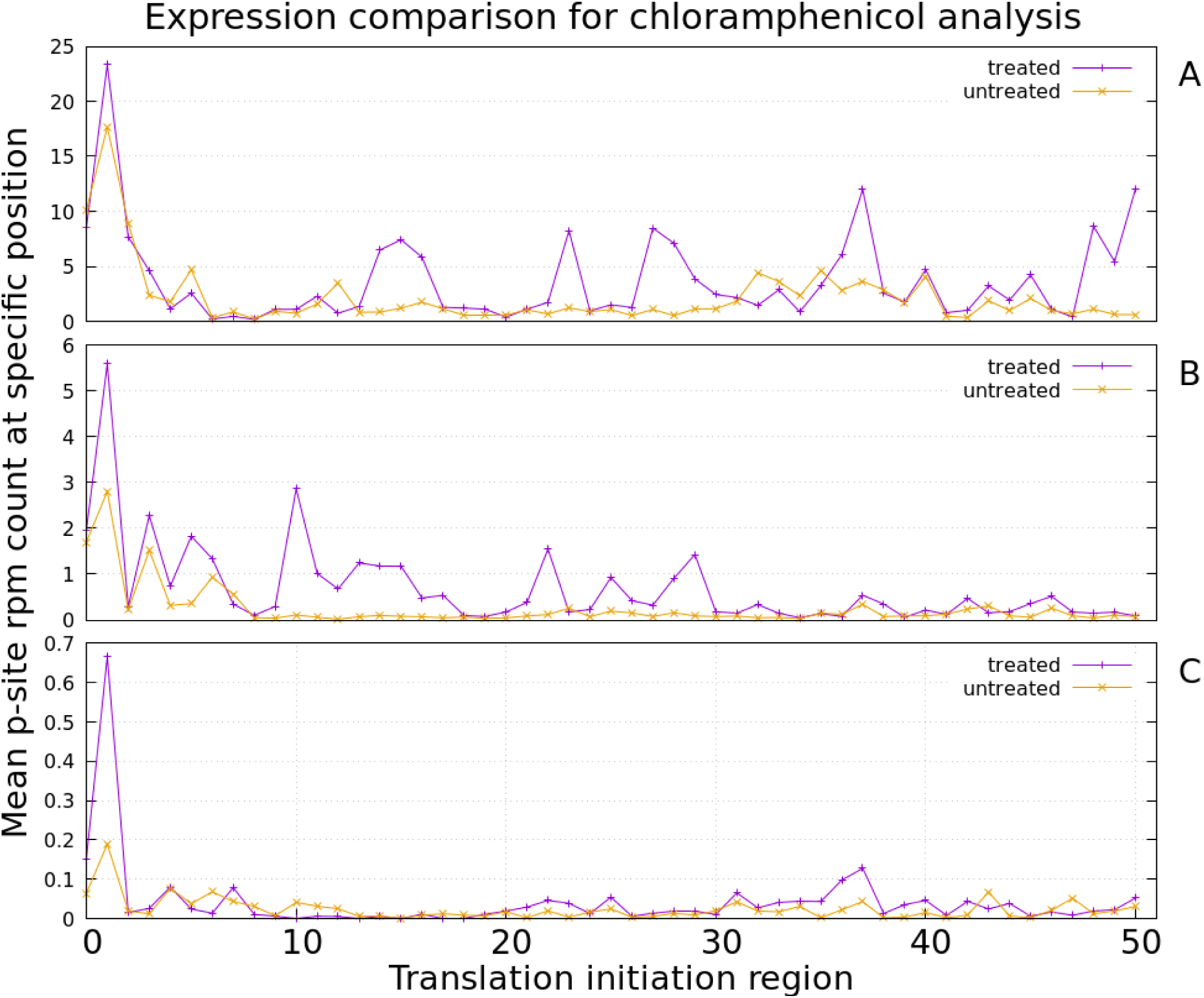
Average read accumulation for all analysed genes being A) highly, B) medium, C) weakly expressed. Chloramphenicol treated samples (purple line) were compared to untreated samples (orange line). The drug treatment in general seems to promote read accumulation at the TIS.

### Sufficient coverage depth for ribosome profiling

Genome coverage is defined as how often a base is sequenced on average during an experiment. Therefore, the total number of reads sequenced determines the overall coverage. In RNA-seq experiments, expression values vary greatly between genes depending on their role in a particular growth stage or environment. To ‘catch’ all genes being transcribed the amount of reads being sequenced needs to be sufficient in order to also detect weakly expressed genes [54]. Increasing the number of reads sequenced beyond a certain point will likely lead to saturation of the set of genes able to be detected. Haas et al. (2012) claimed that a range from 5-10 million reads covering mRNA in RNA-seq approaches is enough to also detect weakly expressed genes while minimising false positive detections. Whether such false positives ever exist could well be discussed. In any case, in RNA-seq, for highly expressed genes an amount of 2 million reads already seems to be sufficient [55]. However, the necessary amount of reads is highly dependent on the characteristics of the experiment. If the particular interest lies in using ribosome profiling experiments to detect weakly expressed genes [28, 30, 56, 57], a certain read number matching a gene is needed to confidently confirm expression (e.g. RPKM above a certain threshold). To detect the necessary read depth, we used an analysis Haas et al. performed in 2012 and adapted the calculation to ribosome profiling data [54]. Prediction of ORFs was conducted as described in the methods section. Figure 6 provides a brief overview.

**Figure 6:**
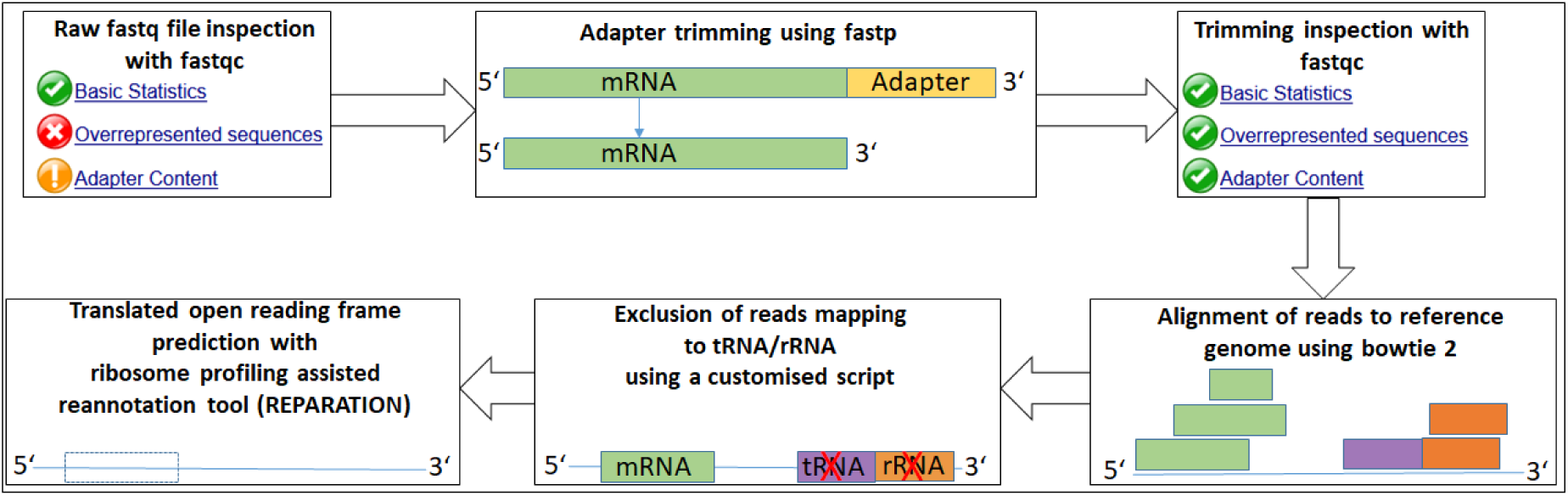
Analysis pipeline for prediction of translated open reading frames.

The total number of reads, excluding the ones mapping to rRNA or tRNA, were compared to the amount of predicted open reading frames by REPARATION representing detectable annotated genes (Figure 7). Haas et al. were able to detect “[…] all but 2 of 4149 ORFs annotated […]” with a threshold of at least one read mapping in an open reading frame for RNA-seq data [54]. Mapping of just a single read possibly is a lenient criterion and thus with our analysis we cannot reach the same detection level. For predictions by REPARATION, we require at least 3 reads mapping to a predicted open reading frame. With this threshold, the number of annotated genes detected in a RIBO-seq experiment reaches saturation with 20 million reads after rRNA/tRNA removal per sample, detecting ~ 3500 genes (82% of 4242 known annotated genes, NC_000913.3). At least 20 million reads mapping to annotated genes seem to be an appropriate amount for ribosome profiling data and does not exceed the amount of reads predicted to be excessive for RNA-seq (50 million) [54]. The consequences for weakly expressed unannotated genes require further research however.

**Figure 7:**
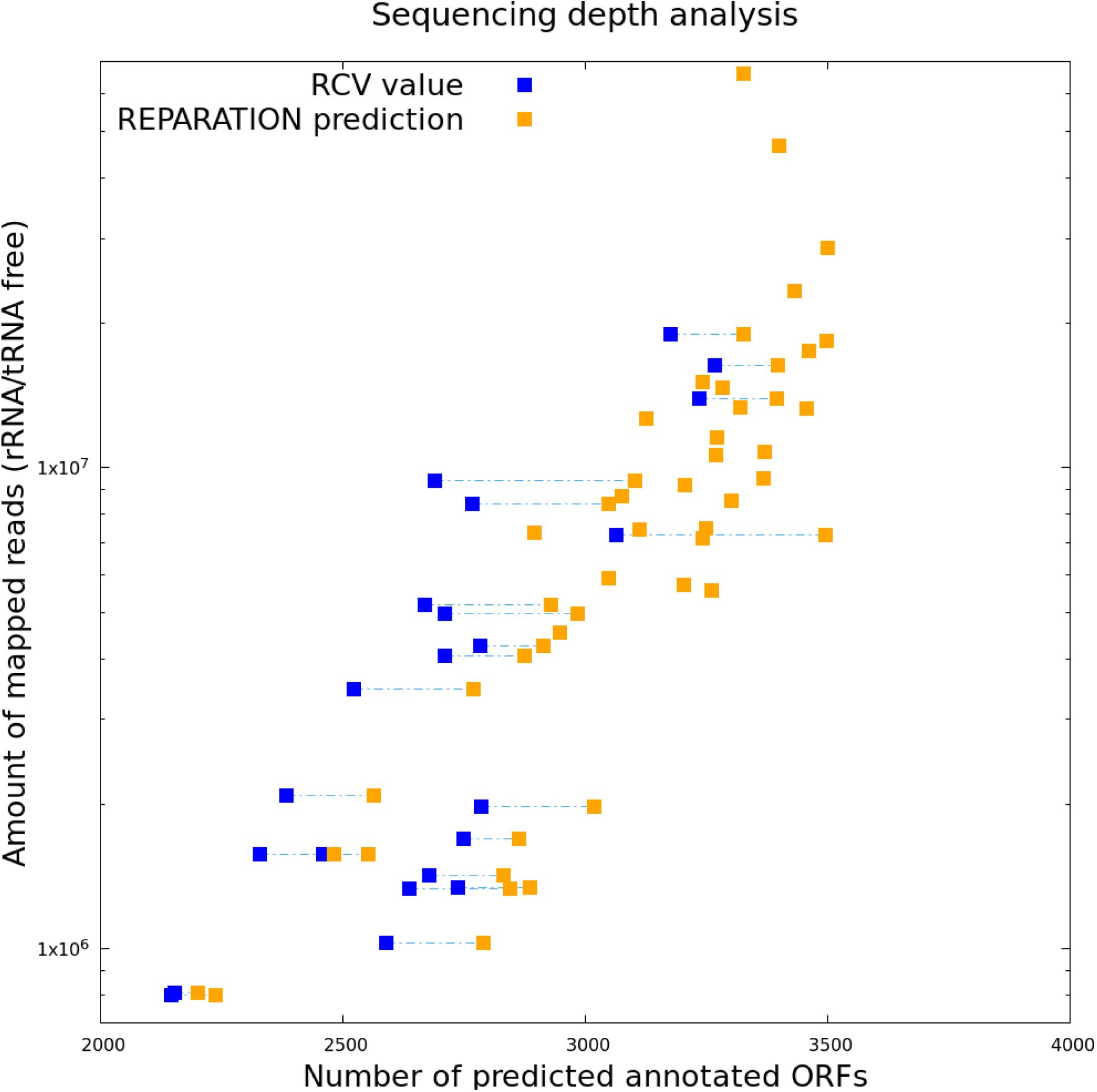
Plot showing relation between amount of reads sequenced (without rRNA/tRNA) and the number of predicted annotated genes. Estimation of annotated genes are based on REPARATION prediction (orange) or RCV value (blue). Samples having both kinds of estimation are connected via dashed lines.

If RNA-seq data was available, RCV values were calculated to obtain a second reference point for the read amount needed in RIBO-seq experiments. For 22 samples, the calculation was performed, showing the distribution in Figure 7. Predictions of genes were considered ‘true’ if having an RCV ≥ 0.355 [27]. For samples for which both a REPARATION based prediction and an RCV value was available, data points were connected with dashed lines in Figure 7. When using an RCV of 0.355 as threshold, fewer genes compared to REPARATION were found. Nevertheless, overall this supports the result that 20 million reads are sufficient to detect most of the annotated genes in RIBO-seq experiments. Extensive further work on this question is however required, with recent improvements in gene prediction from RIBO-seq data [58–60] and many studies on previously unrecognised small proteins to take into consideration [16, 30, 57, 61, 62].

## Discussion

### Size selection

Careful size selection should help to ensure that only ribosome protected footprints are sequenced [5]. In our analysis, we find that a discrepancy between the purported selected size and the actual read length distribution sequenced is common. It has recently been suggested that a range from 15 to 40 nucleotides should be taken into consideration for the experimental size selection and data analysis [6]. We argue that a narrower selection range is beneficial for gene detection, as longer fragments have a higher probability of mapping to either tRNA or rRNA and mapping of shorter reads is more error prone. Using a better size selection, such reads can be excluded from the sample before sequencing, resulting in a higher coverage of protein-coding mRNA reads.

All but one experiment chose ranges for the distribution rather than one specific length. Having a closer look at each distribution and the average trend per experiment, four datasets show an actual distribution within their selection (Bal_14, Hwa_17, Kan_14, Woo_15). The other four sets also had reads of length outside their chosen selection. These results already show that the actual size selection obtained is not only highly dependent on the range chosen but also on the precision with which it is conducted. The accuracy of gel excision techniques is important, and we want to give a recommendation for the area selected to increase RIBO-seq performance. The common range between 20 to 40 nucleotides, should obtain a spectrum with a potential peak value at around 24 nucleotides [6, 31, 45]. Nevertheless, this peak value can also be achieved with a size selection from 20 to 30 nucleotides [38], or even more narrowly, of 23±3 nucleotides for a different *E. coli* strain [23]. We suggest choosing the narrower range, aiming for peak values at around 24-27 nucleotides. These lengths predominantly map to protein-coding mRNA (Figure 3, see below). Interestingly, choosing fragments of 25 to 42 nucleotides in length resulted in a peak value at around 30 nucleotides [24]. It could be argued that shifting the lower boundary value up (around five nucleotides) results in a distribution shift as a whole. The size selection should have little correlation with the fragment length, since this is primarily dependent on the protection by the ribosome from the digestion of nucleases [1, 2, 6]. Possibly, an insufficient performed digestion might have led to longer fragments in this one study (Kan_14).

### Specific read length correspond to different RNA types

rRNA and tRNA make up around 95-97% of the RNA in bacterial cells [54, 63], hence their removal, by either fragment size selection or rRNA depletion before library preparation, would increase the proportion of reads mapping to mRNA. Reads ranging in length from 24 up to 27 nucleotides seem to be the dominant ones for mRNA (Figure 3). tRNA reads appear to be longer than mRNA reads. To exclude excessive tRNAs, size selection should again be below 30 nucleotides. To support the hypothesis of excluding tRNA due to a narrower size selection, the percentage of remaining reads mapping to tRNA was compared (Table S4). The samples with a performed size selection of 20-30 nucleotides contained less than one percent tRNA (Table S4). Also, a size selection spanning from 25 to 31 nucleotides resulted in a remaining tRNA percentage of 11.5 % on average, which is less than with a range from 20-40 for size selection (33,5%). Choosing a cut off for size selection at around 30 nucleotides should ensure exclusion of most of the tRNA. The lowest amounts of reads mapping to rRNA are also found with a performed size selection of 20-30. In contrast, the second lowest remaining rRNA amount (15.2%) can be found with a performed size selection of 25 to 42. Even though the selection is in the higher range, the amount of rRNA is fairly low. These results suggest that for rRNA contamination, a properly performed depletion is as important as size selection. While size selection could exclude longer rRNA and tRNA reads, proper rRNA depletion could additionally minimise the amount of shorter rRNA fragments, increasing the proportion of sequence fragments with protein-coding information.

Reads belonging to rRNA show two peaks at 26 and 31 nucleotides. The longer fragments at 31 nucleotides would be excluded if the size selection threshold did not exceed 30 nucleotides. To exclude remaining rRNA fragments several options for their depletion can be accessed currently. Three experiments specified using commercial kits, namely RiboZero (Illumina; Woo_15, Hwa_17) or MICROBExpress (Thermofisher; Elg_14) for additional rRNA depletion. The remaining experiments performed standard protocol rRNA depletion based on biotin labelled rRNA hybridisation. The average percentage of rRNA remaining in kit-treated samples is not significantly lower than in samples where no commercial kit was used (Table S4). The lowest rRNA amount remaining can be found in a set for which no kit-based depletion method was mentioned (Bal_14). This suggests that size selection might be contributing more effectively to rRNA depletion. However, rRNA fragments with a length of 26 nucleotides cannot be excluded by size selection. Thus, the combination of a narrower selection with a well-performed rRNA depletion is crucial for lower amounts of rRNA present.

### Longer reads in 5’-UTR region

Normally for *E. coli* the main SD sequence is located around eight nucleotides upstream of a start codon and aids in translation initiation due to binding a 16S rRNA of the 30S small subunit, allowing the ribosome to assemble at the coding sequence [64–70]. The interaction of the SD sequence and its counterpart resulting in assembly of the ribosome complex might cause protection of longer fragments than otherwise expected [71, 72]. As already mentioned in the results, if reads with a Shine-Dalgarno like motif are of interest, a range exceeding 22 to 30 nucleotides should be taken into consideration but this will decrease the overall performance.

### Further experimental improvements

To further improve the performance of size selection, another adjustment regarding gel excision can be made. In the early RIBO-seq protocols, a DNA ladder was used as a size comparison [5]. Experiments performed by our group, however, show that using single-strand RNA molecules as a ladder is more suitable. DNA molecules diffuse slower through the gel due to their increased molecular weight size and DNA is less prone to folding. Therefore, orienting gel excision of RNA fragments based on a DNA ladder leads to an incorrect size range and probably larger fragments then intended. We recommend using a single-stranded RNA ladder with a mixture of random sequences of the intended size to ensure a more precise size selection.

### Sufficient coverage depth for ribosome profiling

For RIBO-seq experiments, a higher read amount is necessary to detect annotated genes independent of their expression levels than in RNA-seq experiments. We recommend sequencing to at least 20 million mapped reads (rRNA/tRNA excluded). This should lead to a broad spectrum of strongly and weakly expressed genes detectable.

### Adjustments for detecting new genes

To increase the possible detection of so far unannotated translated ORFs, we want to give a recommendation on what stalling method is most helpful here. Drug induced ribosomal stalling by elongation inhibitors such as chloramphenicol, Onc112, or retapamulin may be the most promising strategy. Due to read accumulation bias at the TIS [6, 16], the start site is more clearly evidenced with these methods – which is very useful when trying to select the correct ORF amongst candidates in the same region. With our analysis, we can confirm the claimed accumulation bias due to chloramphenicol application. This stalling method is likely to therefore aid in detecting undetected genes by improving the resolution of translation start site detection. Additionally, recent studies using retapamulin or Onc112 showed that they are not only suitable in detecting additional translation start sites representing potential length variations of genes [16, 73, 74], but based on the accumulation bias they were also able to distinguish between two genes situated in different reading frames in the same region [16]. With all the named advantages in detecting open reading frames and their exact start site, the use of elongation inhibitors is clearly useful for detecting yet unrecognised translated ORFs.

Undetected ORFs potentially remain in this state partly as they are typically weakly expressed under standard experimental conditions. As all samples used for this analysis were grown in LB medium, the expression of many overlapping genes, for instance, is likely to be low if they typically function in stress response. However, with the accumulation bias, the detection might be improved as it can lead to detecting a new translation start site. The time between adding antibiotics and finally harvesting the bacteria should be kept as short as possible otherwise the translatome is likely to change in response [6]. Some researchers use rapid filtration followed by cell freezing to stall the ribosomes, bypassing the potential change in the translatome due to drugs [6]. However, this read accumulation bias that some want to avoid could also potentially aid in detecting new ORFs.

On the other hand, increasing the sequencing depth could aid in detecting weakly expressed unannotated genes present in a sample. Some studies have claimed that ORFs antisense to annotated genes, detectable by increasing the read depth, mostly result from transcription initiation or termination inaccuracy and therefore are presumably non-functional sequences [54, 75–77]. A number of other researchers have proposed that many antisense RNA sequences are in fact functional [78–80]. We additionally propose the stronger claim that many antisense ORFs with reads mapping to them include undetected protein-coding ORFs. Several studies have shown that there are still unannotated genes present in even the well-studied *E. coli* K12 [81–83]. Phenotypes for recently detected genes overlapping an annotated one (overlapping genes, OLGs) in enterohemorrhagic *E. coli* (EHEC) have been found in different environmental conditions, e.g. salt stress [56], anaerobiosis [28] or arginine supplementation [57] implying that they are present in a relatively low amount when cultivated in standard LB media. Increasing read depth and sampling in other media conditions might improve the ability to predict such low abundance proteins. With predictions based on REPARATION, we had a look at the amount of detected ORFs we classify as being located partially overlapping an annotated gene (pORF) or fully embedded on the alternate strand to an annotated gene (eORF). Surprisingly, for both categories the threshold of 20 million effective reads also seems sufficient (Figure S4A+B) – but the extent to which this is an artefact of the prediction tools used remains to be determined.

## Conclusion

The introduction of ribosome profiling by Ingolia et al. in 2009 was the first stepping stone for directly investigating entire ‘translatomes’, beginning in eukaryotes [2]. Since then, investigation of the bacterial translatome has led to improvements in the experimental protocols. However, there still is no consensus about which size selection to choose for bacteria in order to obtain the most informative ribosomal footprints. Our recommendation is to choose a size selection between 22-30 nucleotides to not only exclude longer fragments associated with tRNAs and rRNAs during size selection but also to enrich for fragments that are highly likely to be protein-coding. However, if researchers are interested in the 5’-UTR region including the Shine-Dalgarno sequence, size selection should be adapted according to the expected fragment length. The total amount of mappable reads required for RIBO-seq appears to be around 20 million effective reads; that is after excluding rRNA/tRNA reads. With this amount of reads most annotated genes that are being translated at the point of harvesting are detectable. However, this result may be limited by the prediction tool used (REPARATION). The RCV threshold provides an additional criterion, which can be used to ensure translation of candidate ORFs. This value is also interesting in studying translation response (e.g., how well certain mRNAs are picked up by ribosomes). Finally, in light of variation in the detail of reporting in existing studies, we emphasize the importance of publishing the complete protocol with all details for future reference and analysis (listed in Table S5). Only with exact information provided, for example concerning ribosome stalling methods, harvesting or size selection, the required comparative analyses can be performed precisely.

