## Supporting information for "Improving Bacterial Ribosome Profiling Data Quality"

Table S1: Overview of experimental methods used in the analysed datasets

| GEO/Experiment number | Stalling method | Harvesting method | Size selection | Publication | Abbreviation |
|---|---|---|---|---|---|
| GSE85540 | - | rapid filtration | 20-40 nt | Hwang et al. 2017 | Hwa_17 |
| GSE68762 | chloramphenicol | centrifugation | n.d. | Bartholomäus et al. 2016 | Bar_16 |
| GSE86536 | chloramphenicol/linezolid | rapid filtration | 28-42 nt | Marks et al. 2016 | Mar_16 |
| E-MTAB-2903 | chloramphenicol | rapid filtration | ~28 nt | Wang et al. 2015 | Wan_15 |
| GSE64488 | - | rapid filtration | 20-40 nt | Woolstenhulme et al. 2015 | Woo_15 |
| SRP048921 | - | rapid filtration | 20-30 nt | Balakrishnan et al. 2014 | Bal_14 |
| SRP040142 | - | rapid filtration | 28-42 nt | Elgamal et al. 2014 | Elg_14 |
| GSE61619 | erythromycin/telithromycin | rapid filtration | 25-42 nt | Kannan et al. 2014 | Kan_14 |
| GSE33671 | chloramphenicol/ - | centrifugation/filtration | 25-31 nt | Oh et al. 2011 | Oh_11 |

Table S2: Softwares used with special settings

| Software | Source | Important settings |
|---|---|---|
| Bedtools v.2.25.0 | Quinlan, A. R. 2014 | Intersect -u; -s; -f 0.55; -sorted<br>Coverage -s; -F 0.5; -sorted |
| Bowtie2 v.2.2.6 | Langmead et al. 2012 | --local; -N 0; -L 19 |
| Fastp 0.14.2 | Chen et al. 2018 | -x; -Q |
| FastQC v0.11.4 | http://www.bioinformatics.babraham.ac.uk/projects/fastqc/ | - |
| Samtools 1.7 | Li et al. 2009 | - |
| REPARATION | Ndah et al 2017. | -sd yes; -min 19 |

Table S3: Expression intensity classification

| Category | High | Medium | Low |
|---|---|---|---|
| RPKM values | 1.000 - 3.000 | 100 - 250 | 10 - 20 |

Table S4: percentage of different RNA types per samples
Values for different types do not add up to 100, remaining reads mapping potentially unannotated ORFs

| Sample | Genes | tRNA | rRNA | Sample | Genes | tRNA | rRNA | Sample | Genes | tRNA | rRNA |
|---|---|---|---|---|---|---|---|---|---|---|---|
| SRR1734437 | 29.04 | 23.35 | 45.42 | SRR4023280 | 27.94 | 22.3 | 44.04 | SRR1200739 | 20.49 | 35.33 | 41.84 |
| SRR1734438 | 35.16 | 27.89 | 34.57 | SRR1613263 | 81.76 | 0.41 | 14.64 | SRR1200750 | 16.36 | 34.66 | 46.52 |
| SRR1734439 | 52.05 | 16.59 | 29.42 | SRR1613265 | 82.41 | 0.44 | 13.89 | SRR1200751 | 18.35 | 39.41 | 41.13 |
| SRR1734440 | 9.54 | 68.82 | 20.34 | SRR1613266 | 77.73 | 0.39 | 18.84 | SRR1583082 | 61.92 | 23.19 | 11.01 |

| | | | | | | | | | | | |
|---|---|---|---|---|---|---|---|---|---|---|---|
| SRR1734441 | 42.12 | 17.63 | 37.69 | SRR1613268 | 84.34 | 0.42 | 11.54 | SRR1583083 | 53.8 | 23.75 | 17.38 |
| SRR1734442 | 41.29 | 14.97 | 41.8 | SRR1613269 | 35.3 | 0.18 | 63.2 | SRR1583084 | 58.44 | 21.15 | 17.24 |
| SRR1734443 | 36.6 | 18.05 | 43.01 | SRR1613270 | 83.64 | 0.39 | 12.41 | SRR4190324 | 33.84 | 12.72 | 50.49 |
| SRR1734444 | 48.09 | 15.37 | 34.89 | SRR1613272 | 84 | 0.42 | 12.01 | SRR4190325 | 43.07 | 19.28 | 32.08 |
| SRR4023274 | 59.17 | 16.6 | 22.73 | SRR1613277 | 72.1 | 0.72 | 13.03 | SRR4190326 | 28.46 | 15.25 | 51.71 |
| SRR1613278 | 79.97 | 0.68 | 16.55 | SRR1200730 | 25.13 | 33.65 | 38.45 | SRR364363 | 51.95 | 7.93 | 36.79 |
| SRR1613280 | 82.73 | 0.67 | 13.5 | SRR1200731 | 18.09 | 29.53 | 50.38 | SRR364364 | 45.11 | 18.2 | 32.11 |
| SRR1613281 | 78.78 | 0.62 | 17.9 | SRR1200738 | 25.84 | 18.61 | 52.64 | SRR364365 | 55.6 | 8.37 | 30.36 |
| SRR1613283 | 85.17 | 0.7 | 10.95 | SRR2016457 | 42.69 | 10.83 | 40.66 | SRR364366 | 48.46 | 19.03 | 27.52 |
| SRR1613285 | 61.99 | 0.48 | 35.39 | SRR2016465 | 29.75 | 1.47 | 66.06 | SRR364367 | 54.2 | 9.2 | 33.84 |
| SRR1613287 | 73.35 | 0.36 | 23.54 | SRR364370 | 62.91 | 12.15 | 20.76 | SRR364368 | 63.72 | 11.34 | 19.7 |
| | | | | | | | | SRR364369 | 38.67 | 5.81 | 53.95 |

Table S5: Minimum experimental information which should be reported.

| | |
|---|---|
| Adapter sequence | Digestion enzyme + concentration |
| Medium supplementation | Polysome fraction |
| Stalling method | Size selection range |
| Harvesting method | Depletion treatment |

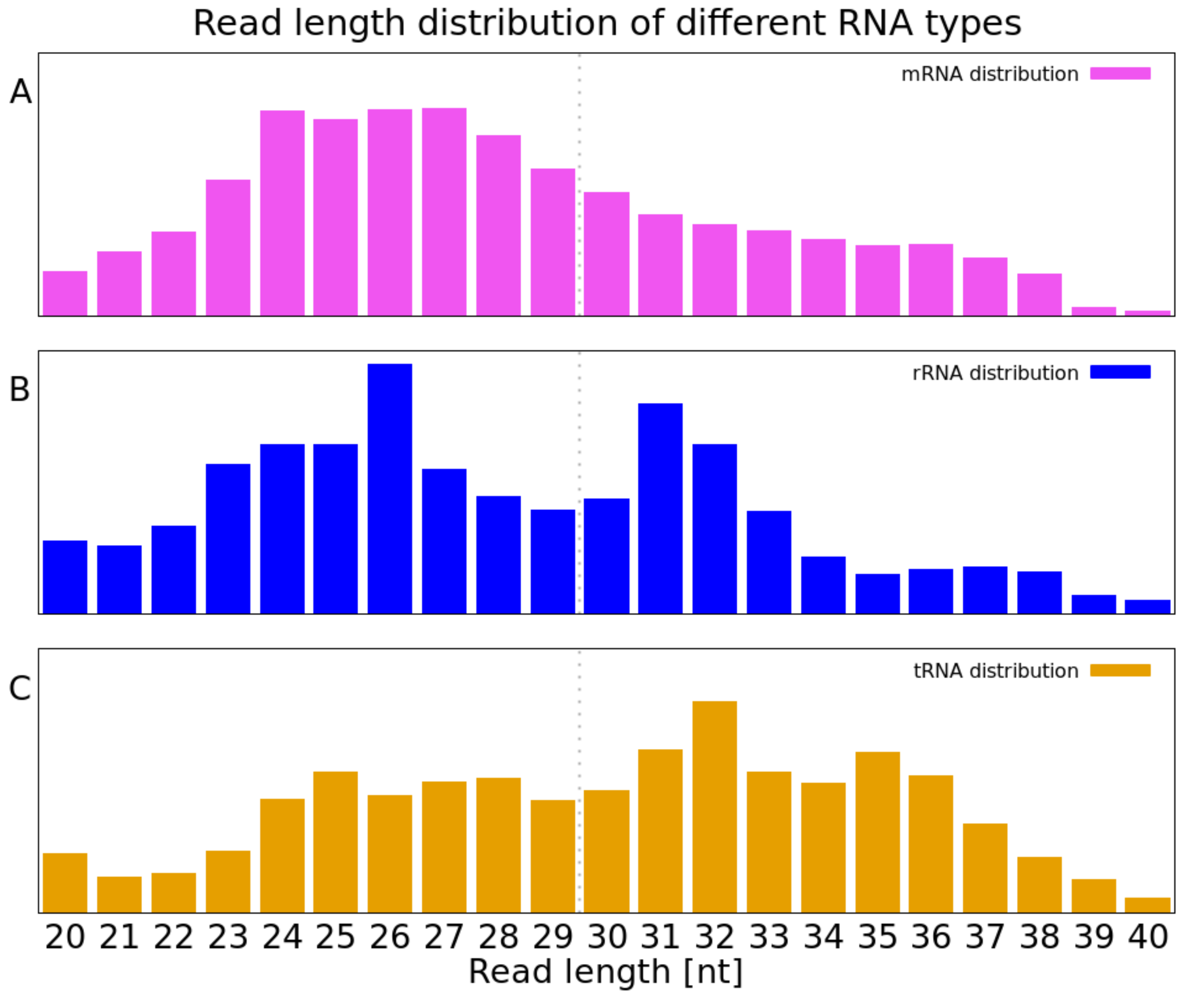


Figure S1: Analysis of mean value for length distribution within one RNA category (n=46), amount of reads at a specific length was used to calculate mean out of the amount of all mapping reads within the RNA type:
A) Distribution of reads mapping only to mRNA; B) Distribution of reads mapping only to rRNA; C) Distribution of reads mapping only to tRNA; dotted line showing suggested size selection border

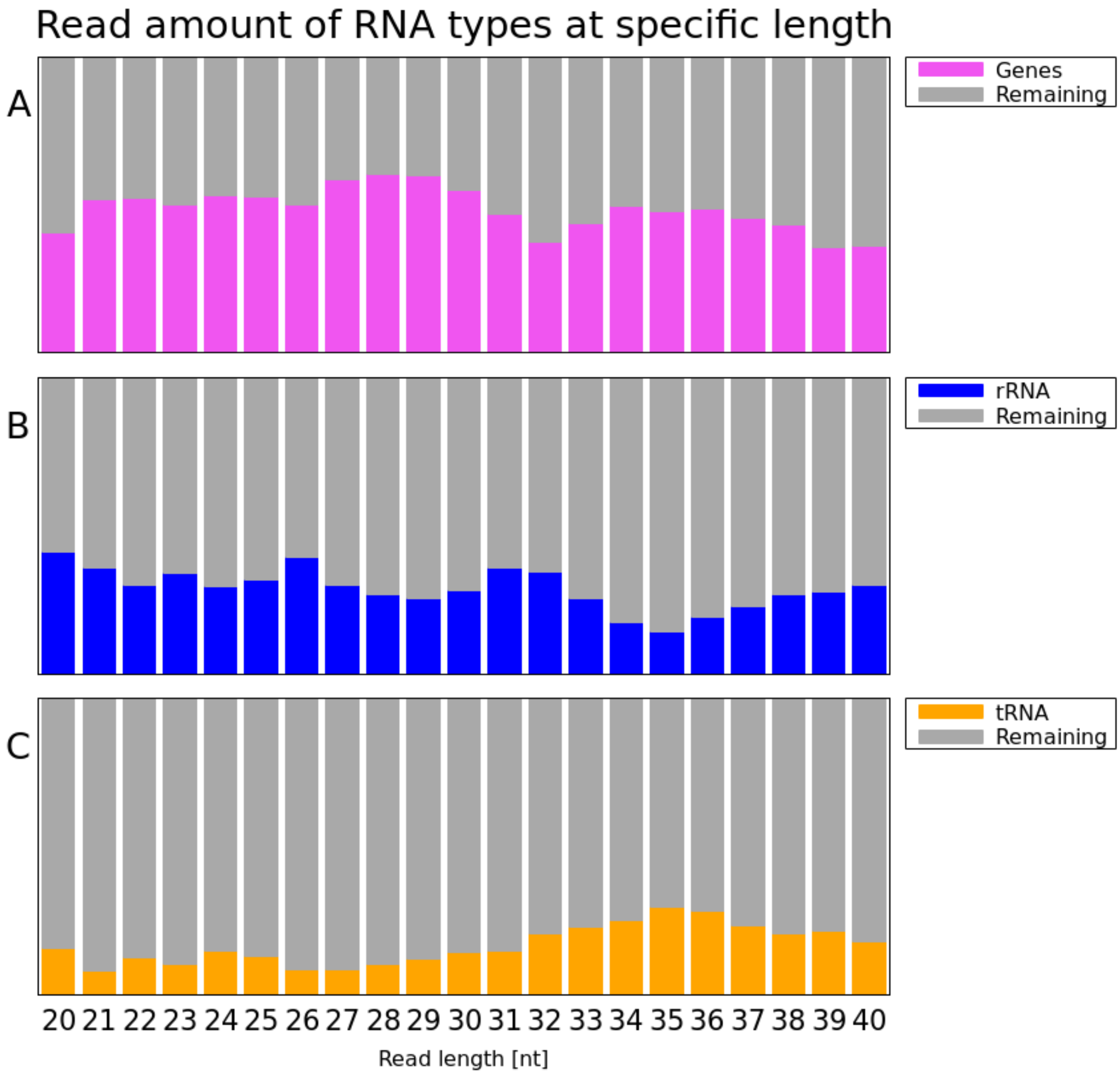


Figure S2: Analysis of mean values for different RNA types at a specific read length; A) Distribution of reads mapping only to mRNA; B) Distribution of reads mapping only to rRNA; C) Distribution of reads mapping only to tRNA

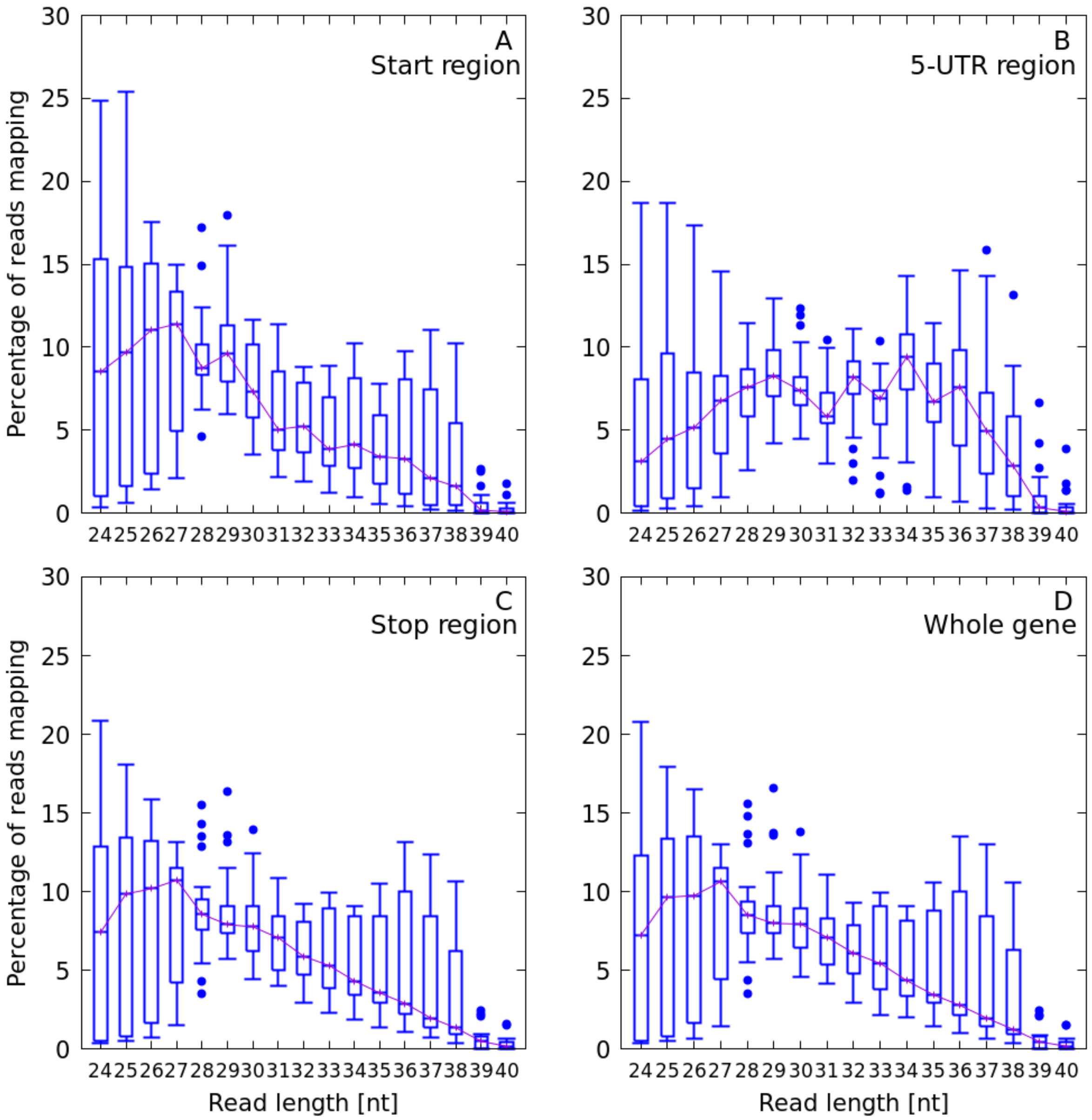

Figure S3: Read length of reads mapping in A) the start region; B) in 5'-UTR; C) the stop region; D) over the whole gene

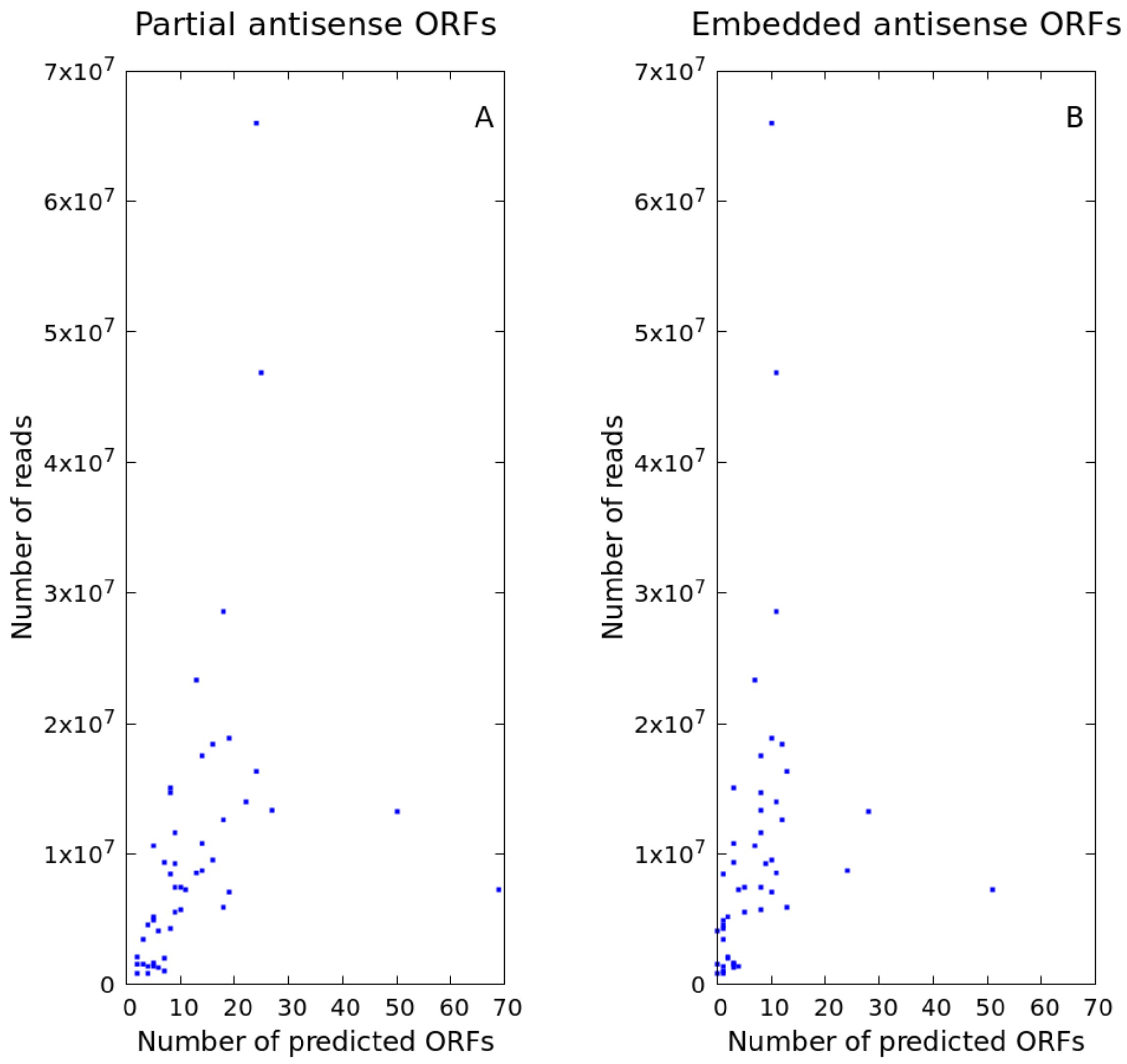


Figure S4: Sequencing depth analysis for A) partial antisense overlaps B) embedded antisense ORFs
